# GenFam: A web application and database for gene family-based classification and functional enrichment analysis

**DOI:** 10.1101/272187

**Authors:** Renesh Bedre, Kranthi Mandadi

**Author notes:** Correspondence: Kranthi Mandadi.

## Abstract

Genome-scale studies using high-throughput sequencing (HTS) technologies generate substantial lists of differentially expressed genes under different experimental conditions. These gene lists need to be further mined to narrow down biologically relevant genes and associated functions in order to guide downstream functional genetic analyses. A popular approach is to determine statistically overrepresented genes in a user-defined list through enrichment analysis tools, which rely on functional annotations of genes based on Gene Ontology (GO) terms. Here, we propose a new approach, GenFam, which allows classification and enrichment of genes based on their gene family, thus simplifying identification of candidate gene families and associated genes that may be relevant to the query. GenFam and its integrated database comprises of three-hundred and eighty-four unique gene families and supports gene family classification and enrichment analyses for sixty plant genomes. Four comparative case studies with plant species belonging to different clades and families were performed using GenFam which demonstrated its robustness and comprehensiveness over preexisting functional enrichment tools. To make it readily accessible for plant biologists, GenFam is available as a web-based application where users can input gene IDs and export enrichment results in both tabular and graphical formats. Users can also customize analysis parameters by choosing from the various statistical enrichment tests and multiple testing correction methods. Additionally, the web-based application, source code and database are freely available to use and download. Website: http://mandadilab.webfactional.com/home/. Source code and database: http://mandadilab.webfactional.com/home/dload/.

## INTRODUCTION

In recent years, genome-wide analyses using high-throughput sequencing (HTS) technologies, have become indispensable to life science research. Generating large-scale datasets has become relatively straightforward, as opposed to efficiently interpreting the data to gain intuition into biologically significant mechanisms. Data mining tools that determine, predict, and enrich putative functions among HTS datasets are highly valuable for such genomic analyses (Backes et al., 2007). For instance, RNA-sequencing (RNA-seq) analysis is a high-throughput approach to study transcriptome regulation by determining transcript-level changes in multiple cell- or tissue-types, or among varying experimental conditions (e.g., unstressed vs. stressed). In a typical RNA-seq experiment, the analysis yields hundreds, if not thousands, of genes that are differentially expressed among the experimental conditions. Uncovering enriched biological pathways among these gene lists is a valuable starting step for downstream functional genetic analyses.

The Gene Ontology (GO)-term based enrichment tools (e.g., BinGO (Maere et al., 2005), Blast2GO (Conesa et al., 2005), AgriGO (Du et al., 2010), PlantGSEA (Yi et al., 2013)) are widely used by researchers to infer the biological mechanisms of genes identified in HTS experiments (Mandadi and Scholthof, 2012; Chen et al., 2013; Bedre et al., 2015; Mandadi and Scholthof, 2015; Bedre et al., 2016; Li et al., 2017; Bedre et al., 2019). These tools identify overrepresented GO terms associated within a user-defined list of genes by mapping them to the background genome annotations and calculating statistical probability of the enrichment relative to the background. The enrichment tools can classify genes into GO categories or pathways related to biological process, molecular function and cellular locations (Goffard and Weiller, 2007; Du et al., 2010). The GO-enrichment and the resultant hierarchy are very useful to understand the complex biological processes that are being enriched. However, information on specific biological attributes of a gene, such as the gene family (a group of homologous genes with common evolutionary origin and biological functions) level information, are hard to glean from GO-enrichment alone (Ashburner et al., 2000; Lee et al., 2005). For instance, enrichment of a transcription factor will fetch GO terms for “regulation of transcription (GO:0006355)” or “DNA binding (GO:0003700)” or “response to stress (GO:0006950)” but does not identify which transcription factor family genes (e.g., WRKY, bZIP) being enriched. Having this information, allows users to readily interpret large-scale datasets effectively and select favorite gene families for further functional studies. While providing the information for functional studies, gene families also could reveal the accurate gene annotation information that could not be easily determined by BLAST-based tools alone. Further, comparative gene family size analysis can certainly be informative and valuable approach to explore the biologically relevant functions related to genome architecture and adaptation or speciation of various plant species (Guo, 2013).

With the availability of complete genomes and sequence data, identification, and analysis of specific gene families among plant species has become necessary. In this study, we present a unique approach to perform classification and enrichment of genes to identify overrepresented gene families (GenFam) in a user-defined query list. We suggest that GenFam is a valuable addition to a plant biologists toolkit to analyze large-scale HTS datasets. By determining overrepresented gene families in a user-defined gene list, rather than GO terms or hierarchy alone, GenFam empowers users to readily interpret information of gene families (e.g. WRKY, bZIP) in their queries, and move forward to selecting favorite overrepresented genes (or families) for downstream studies and interpretation. GenFam is also freely accessible to users on the world-wide web, as a user-friendly, graphical-user interface.

## MATERIALS AND METHODS

### Background database

GenFam currently supports the analysis of sixty plant genomes. GenFam classifies genes into 384 representative and unique gene families, which to the best of our knowledge the largest collection, based on the well-annotated *Arabidopsis thaliana* (Berardini et al., 2015) and rice (*Oryza sativa*) (Kawahara et al., 2013) genomes, literature search, and Pfam protein families database (El-Gebali et al., 2019). We have identified and used Pfam common conserved domains and domain organization among the homologous gene sequences to assign the gene families. These highly conserved domains define protein functions and classifies protein-coding genes into gene families. The conserved signature protein domains have the ability to detect the divergent or distantly related homologs which would be prohibitive with sequence based similarity analysis tools [e.g. BLAST (Altschul et al., 1997)]. Therefore, domain-based search method would identify more genes belonging to gene families than BLAST-based homology search.

To identify and classify gene families in plants, we have leveraged the publicly available genomic resources at Phytozome (v12) database. The protein sequences of sixty plant genomes were used to identify conserved protein domains to assign families to known and unclassified or novel genes. The respective protein domains were predicted by HMMER (v3.1b2) using a protein family hidden Markov model (HMM) profiles (Pfam release 32.0) (El-Gebali et al., 2019). We have established rules to classify and assign the genes to gene families based on the presence of signature conserved protein domains and have provided in **Supplementary Table S1**. This approach allowed us to maximize classification including orphan genes with missing annotations, genes with incorrect annotations, and novel genes present among the respective genome databases. Lastly, the background databases were curated to remove redundancy and duplication of gene members among families. In summary, we were able to integrate 384 representative gene families and corresponding (on an average ~41%) genes from sixty plant genomes into our database (**Supplementary Table S2)**. This is a the most comprehensive and largest collection of gene families spanning sixty plant species, when compared to other existing databases. For instance, the recently published gene family database in poplar (GFDP) has classified 6551 poplar genes into 145 gene families derived from Arabidopsis genome (Wang et al., 2018). PlantTFDB (v4.0) and PlnTFDB (v3.0) has classified the genes into 58 and 84 transcription factor gene families (Perez-Rodriguez et al., 2010; Jin et al., 2017). Similarly, another database and analysis toolkit, PlantGSEA, supports the gene family analysis for 13 plant species which mostly imports gene families from well-annotated genomes such as rice (118 gene families) and maize (81 gene families) (Yi et al., 2013).

All the gene family data was formatted using the PostgreSQL database to perform classification and enrichment analysis using various statistical enrichment methods. The GenFam database with complete protein domain annotation and gene family classification can be downloaded from the GenFam website (http://mandadilab.webfactional.com/home/dload/). Detailed statistics for the number of genes assigned to each gene family and the total number of background genes are provided in **Supplementary Table S2**.

### Statistical enrichment methods

GenFam performs three main functions: i) Annotation ii) classification, and iii) enrichment of a user-defined gene list to provide gene family-level attributes. The enrichment is based on the singular enrichment analysis (SEA) method, which computes enrichment of a user-defined list of genes with a precomputed background dataset (Huang da et al., 2009). GenFam accepts different types of gene IDs for the analysis. For example, for rice, it accepts gene (e.g., LOC_Os01g06882) and transcript (e.g., LOC_Os01g06882.1) IDs from parent database such as the Rice Genome Annotation Project (http://rice.plantbiology.msu.edu/). Additionally, GenFam also accepts Phytozome PAC IDs for a given gene (e.g., 24120792 for LOC_Os01g06882), which provides additional flexibility in performing the analysis. To determine an acceptable ID, the user can run the “check allowed ID type for each species” function on the GenFam analysis page (http://mandadilab.webfactional.com/family/). Once the appropriate gene IDs are provided, GenFam classifies and identifies specific gene families and members that are overrepresented in the input gene list.

Even though there is no defined standard for choosing a reference background, it is ideal to select a background that will increase coverage (or intersection) with an input gene list, as well as that enhances specificity of the enrichment analysis (Huang da et al., 2009). GenFam utilizes the number of total genes categorized/annotated into gene families in each plant species as a reference background, rather than using the whole genome. This feature greatly improves the specificity of the enrichment analysis by implementing statistically stringent criteria. For instance, for case study 1, if enrichment analysis was performed with the whole genome as background, it would result in 35 enriched gene families with much lower P-values, when compared to using the current GenFam background (29 enriched gene families) (**Supplementary Table S3**).

GenFam can employ standard statistical tests such as the Fisher exact, Chi-Square (*χ*^2^), Binomial distribution and Hypergeometric tests for enrichment, along with multiple testing corrections to control a false discovery. We recommend using Fisher exact, Chi-square (*χ*^2^) and χ Hypergeometric tests for smaller datasets (<1000) (McDonald, 2009), and Binomial distribution for larger datasets (Khatri and Draghici, 2005; Zheng and Wang, 2008). Furthermore, the Chi-Square (*χ*^2^) test would be appropriate when the user defined gene list has less overlap with the χ background dataset. As a default test, GenFam performs the Fisher exact test, which relies on the proportion of observed data, instead of a value of a test statistic to estimate the probability of genes of interest corresponding to a specific category.

To address the false positives resulting from multiple comparisons especially when the input gene list is large (>1000), GenFam subsequently employs false discovery correction methods including the Benjamini-Hochberg (Benjamini and Hochberg, 1995), Bonferroni (Bonferroni, 1936) and Bonferroni-Holm (Holm, 1979). The various statistical tests and false discovery correction methods can be customized by the user as appropriate.

### Output summary

A snapshot of the analysis page and workflow is shown in Figure 1. Users have the option to either use the default settings or select desired statistical parameters. The analysis page also guides the users to select gene IDs that are acceptable in GenFam (Figure 1). Users are directed to the results after analysis is completed (Figure 1). The results of GenFam analysis are displayed as summary table (HTML) and graphical chart plotted using the -log_10_(P-Value) scores. Higher the -log_10_(P-Value) value, greater the confidence in enrichment of the gene family (Figure 2). The enriched and non-enriched gene family results can also be downloaded as tabular files, with further details of associated P-value and FDR statistics, gene family size, gene IDs and GO terms.

**Figure 1.**
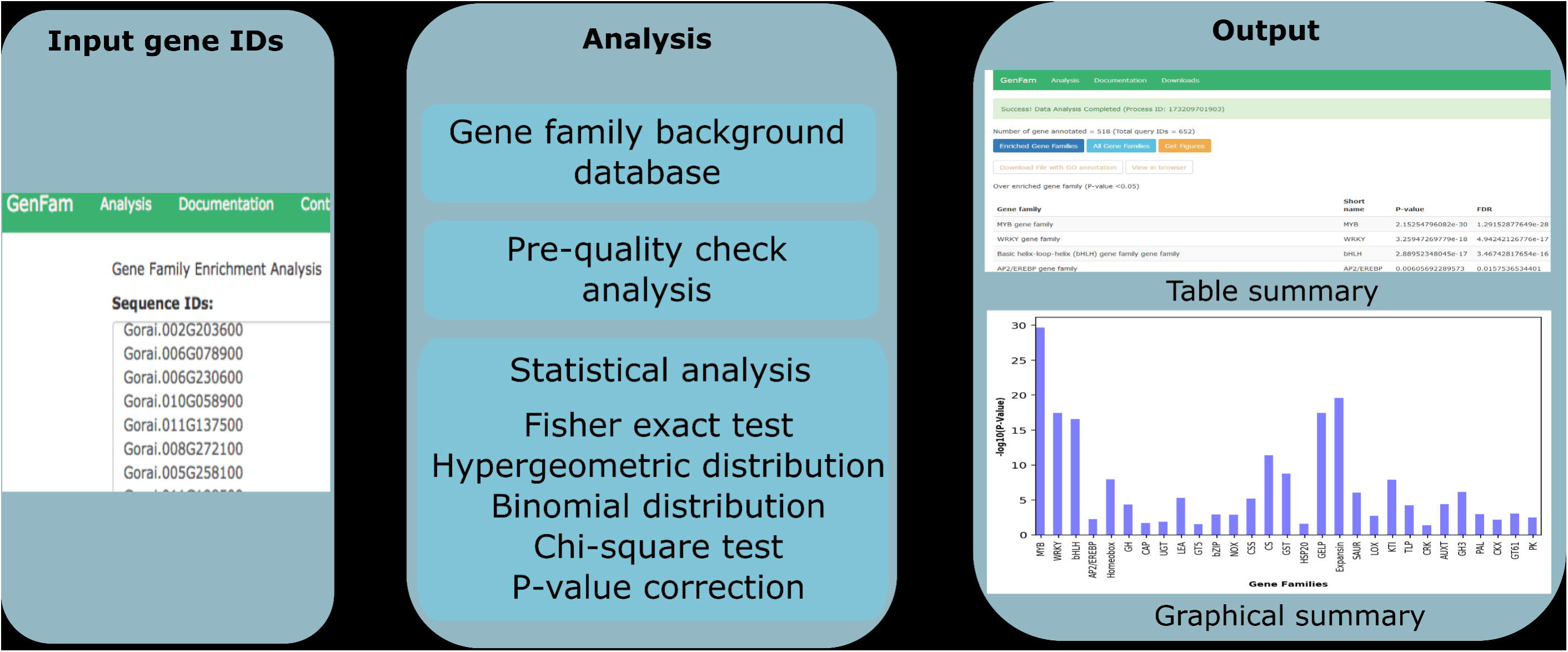
GenFam workflow. The list of input gene IDs for respective plant species provided by the user are analyzed for enrichment analysis using various statistical tests. The ouput of the analysis can be viewed and/or downloaded as a table and/or graphical summary. The results page has multiple options to visualize or download data for both enriched and non-enriched categories (all gene families). The detailed output data from case studies are provided in **Supplementary Tables S3**, **S4**, **S5** and **S6**.

**Figure 2.**
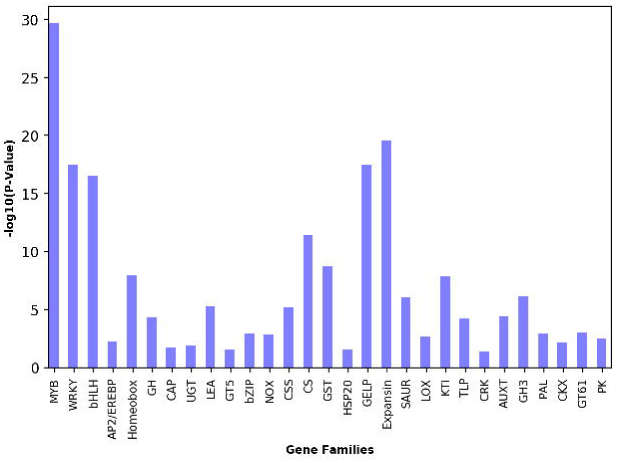
Graphical summary of GenFam enrichment analysis of a cotton case study. Results are plotted as bar chart using the -log_10_(P-Value) scores. Higher the -log_10_(P-Value) value, greater the confidence in enrichment of the gene family.

Along with enrichment results for the gene families, GenFam also provides information related to GO terms in biological process, molecular function and cellular component categories associated with the enriched gene families. In addition to GO terms, GenFam also provides the gene family size and gene IDs associated with each gene family. These results can be downloaded as a tabular file (“Enriched Families”) or as a graphical figure of the enriched families (“Get Figures”). If users only want to retrieve the classification of genes, GenFam parses another tabular file containing the information of all annotated gene families (“All Families”).

### Web server implementation

The GenFam web server is implemented using Python 3 (https://www.python.org/), Django 1.11.7 (https://www.djangoproject.com/) and PostgreSQL (https://www.postgresql.org/) database. All the codes for data formatting and statistical analysis are implemented using Python scripting language. Python is a fully-fledged programming language which offers well developed packages for statistical analysis, graphics and integration with web apps. Therefore, we have chosen Python over other languages such as R for development of GenFam. The high-level Python web framework was constructed using Django. The Python web framework was hosted using WebFaction (https://www.webfaction.com/). The web-based templates were designed using Bootstrap, HTML, and CSS. The GenFam is compatible with all major browsers including Internet Explorer, Microsoft Edge, Google Chrome, Mozilla and Safari. All the precomputed plant gene family background databases were built using advanced PostgreSQL database. The analyzed data was visualized using the matplotlib (Droettboom et al., 2016) Python plotting library.

## RESULTS AND DISCUSSION

### Case studies and analysis

To demonstrate the utility of GenFam, we performed four case studies using transcriptome datasets related to plants from different clades and families (cotton, tomato, soybean and rice) (Bedre et al., 2015; Dametto et al., 2015; Zeng et al., 2017; Cui et al., 2018). We have previously identified 662 differentially expressed genes in cotton (*Gossipium raimondii*, family Malvaceae) infected with *Aspergillus flavus* (Bedre et al., 2015). For the first case study, we used GenFam to determine the enriched gene families among these differentially expressed genes, using the options of Fisher exact test for statistical enrichment, and the Benjamini-Hochberg (Benjamini and Hochberg, 1995) method to control false discovery rate (FDR). Among the 662 genes, 514 genes were annotated and classified into gene families, resulting in ~78% intersection/coverage with the GenFam database. The GenFam enrichment analysis revealed overrepresented gene families such as expansins, kinases, reactive oxygen species (ROS) scavenging enzymes, defense related genes, heat shock proteins and transcription factors—genes that we have hypothesized to mediate cell-wall modifications, antioxidant activity and defense signaling in response to *A. flavus* infection (Bedre et al., 2015). Additionally, GenFam also identified new enriched gene families such as bHLH, GH3, glycosyltransferases and thaumatin that were not reported or identified (Figures 1 and 2; **Supplementary Table S3)**. In the second case study, we analyzed 758 genes which were up-regulated in a cold-tolerant rice (*Oryza sativa*, family Poaceae) (Dametto et al., 2015). Among the 758 genes, 460 genes were annotated and classified into gene families by GenFam, resulting in ~61% intersection/coverage with the GenFam database. GenFam was able to successfully determine enriched gene families related to aquaporins, glutathione S-transferases (GST), transporters, lipid metabolism, transcription factors as well as gene families involved in cell wall-related mechanisms **(Supplementary Table S4)** —genes that were hypothesized by Dametto *et al*. (2015) (Dametto et al., 2015) to play a role in the rice cold stress response. Additionally, GenFam also identified new enriched gene families such as aldehyde dehydrogenase (ADH), kinesins, glycosyltransferases, tubulin, phenylalanine ammonia lyase (PAL) and thaumatin that were not reported or identified **(Supplementary Table S4)**. Next, we analyzed the differentially regulated genes from tomato (*Solanum lycopersicum*, family Solanaceae) (Cui et al., 2018) and soybean (*Glycine max*, family Fabaceae) (Zeng et al., 2017) using GenFam (**Supplementary Table S5 and S6**). We obtained ~65% and ~59% intersection/coverage with the GenFam database for tomato and soybean respectively. The GenFam results in both these studies revealed enrichment of several gene families that were overrepresented and reported by Cui *et al*. (2018) (Cui et al., 2018) and Zeng *et al*. (2017) (Zeng et al., 2017) (**Supplementary Table S5 and S6)**. Additionally, GenFam also identified new enriched gene families such as aquaporins, VQ, tify, GST, and PAL in tomato, and BET, Dirigent, Expansins, Asparagine synthase (ASNS), and Carbonic anhydrase (CA) in soybean that were not reported or identified (**Supplementary Table S5 and S6**). The detailed statistics of enriched gene families for these case studies are provided in **Supplementary Table S3**, **S4**, **S5** and **S6**.

### GenFam advantages and comparison with preexisting enrichment tools

To the best of our knowledge, there is only one existing enrichment tool that comes close to the GenFam approach, i.e., PlantGSEA (Yi et al., 2013), which also allows users to enrich gene lists using gene family attributes. Hence, we performed a comparative analysis of GenFam and PlantGSEA with a dataset from cotton (662 genes)(Bedre et al., 2015) and employing identical parameters (Fisher exact test and Benjamini-Hochberg method) for enrichment. GenFam enriched gene families belonging to cell-wall modifying genes, ROS scavenging genes, transcription factors, lipid metabolism, and stress responsive gene families, both new and previously shown to be biologically-relevant during *A. flavus* infection of cotton (Bedre et al., 2015), while PlantGSEA missed several of these categories **(Supplementary Table S3 and S7)**. Upon further examination, we found that several gene family categories such as the ABC transporters, expansins, and glutathione-S-transferase were absent in the PlantGSEA *G. raimondii* background database. Moreover, PlantGSEA supports only thirteen plant genomes with several redundant and overlapping genes and gene families, which could impact the accuracy of the enrichment analysis. For instance, in the *A. thaliana* genome there are 37 annotated “C2-C2 Dof” transcription factors. PlantGSEA categorized 36 out of the 37 genes into a “C2-C2 Dof” family, but also into an additional “Dof” family leading to redundant gene family categories. GenFam avoids such discrepancies by curation and filtering redundant categories.

Taken together, we suggest that GenFam is a comprehensive and robust gene family classification and enrichment program over prevailing tools, with several advantages: i) GenFam is a dedicated and comprehensive platform for gene family-level classification, annotation and enrichment analysis and supports sixty plant genomes including model and non-model plant species. ii) GenFam background dataset was constructed from well-annotated gene families of *A. thaliana* and rice genomes, literature search, and as well as a systematic HMM profile search for signature conserved protein domain analysis using the Pfam database. This inclusive strategy enabled us to categorize most of the genes into families, including those which may lack a defined annotation in their corresponding genome database or could be novel genes. As a result, GenFam database is by far the largest collection of gene families (384 families). In contrast, existing databases such as PlantGSEA and GFDP only relies on annotations defined by other databases such as TAIR and MSU annotations and/or other transcription factor databases (Yi et al., 2013; Wang et al., 2018). The lack of additional analysis of protein domains perhaps explains the poor representation of gene families in PlantGSEA and GFDP databases. iii) GenFam background dataset was curated to remove redundancy and overlapping genes into different gene families, that enhances the accuracy of the analysis. iv) In contrast to PlantGSEA, GenFam uses the annotated gene families as reference background instead of the whole genome. This feature ensures decreasing enrichment bias and increasing the accuracy of the analysis (Huang da et al., 2009). v) GenFam accepts multiple input IDs including, gene IDs, transcript IDs and PAC IDs, however PlantGSEA and GFDP are restricted to using only gene IDs. vi) GenFam can be solely used for gene family annotation and classification regardless of enrichment analysis if a user is only interested in annotating genes.

## CONCLUSION

Data mining of big datasets (e.g., HTS data) is a very important step, and approaches that can systematically mine biologically relevant information from big data are highly desirable. GO term-based enrichment analyses, although very useful to gain insight about the complex biological information, does not reveal specific gene family level attributes or overrepresented gene families. GenFam can be used as a complementary or alternative approach to GO-based enrichment to interpret biologically relevant information in big datasets by classifying and enriching gene families within a user-defined gene list. This specific information on which gene families are overrepresented allows users to readily identify favorite genes for downstream inquiries. Along with enriching gene families, GenFam can be useful to annotate the large list of genes generated from HTS experiments irrespective of enrichment analysis. In conclusion, we suggest that GenFam would be a valuable and powerful tool for plant biologists utilizing genomics strategies to study plant biology and functional genetics.

## AVAILABILITY AND REQUIREMENTS

**Project name:** GenFam

**Project home page:** http://mandadilab.webfactional.com/home/

**Operating system(s):** Platform independent

**Programming language:** Python 3, Django 1.11.7

**License:** CC BY-NC-ND 4.0

**Any restrictions to use by non-academics:** License needed

## CONFLICT OF INTERESTS

The authors declare no competing financial interests.

## AUTHOR CONTRIBUTIONS

RB conceived the project, developed the database/webserver, performed the case studies and prepared the manuscript. KKM supervised the study, data analysis and interpretation. Both authors have read, reviewed and approved the manuscript.

## ACKNOWLEDGEMENTS

We thank Sonia Irigoyen (Texas A&M AgriLife Research) for review and comments during the preparation of this manuscript. All experiments were conducted following the guidelines and appropriate permissions of the Institutional Biosafety Committee of Texas A&M University. This work was supported by funds from Texas A&M AgriLife Research Insect-vectored Disease Seed Grant to KKM.

## SUPPLEMENTARY MATERIAL

**Supplementary Table S1:** The classification of gene families and assignment of conserved protein domain to each gene family

**Supplementary Table S2:** GenFam database statistics for total number of genes classified into gene families and background number of genes in each plant species

**Supplementary Table S3:** List of the differentially regulated genes and analysis output of the cotton case study

**Supplementary Table S4:** List of the differentially regulated genes and analysis output of the rice case study

**Supplementary Table S5:** List of the differentially regulated genes and analysis output of the tomato case study

**Supplementary Table S6:** List of the differentially regulated genes and analysis output of the soybean case study

**Supplementary Table S7:** PlantGSEA result for gene family enrichment analysis using *G. raimondii* dataset used in GenFam case study.

## Notes

#### Summary of Updates

The manuscript was revised to include four case studies (two new) and a more comprehensive revision of the database and statistics are provided. Additional formatting was performed to enhance clarity and flow of the sections.

